# How Healthy Older Adults Walk on Winding Paths with Different Visual Contrasts

**DOI:** 10.64898/2026.11.13.765168

**Authors:** Sarah E. Overby, Carlie J. Hernjak, Tarkeshwar Singh, Jonathan B. Dingwell

## Abstract

One in four older (age >65 years) adults falls each year, precipitating many hip fractures and traumatic brain injuries. These falls commonly occur during walking, specifically when weight shifting in the lateral direction. Humans must make successful lateral maneuvers in everyday walking tasks, like navigating from room-to-room in their home. Visual contrast sensitivity also declines with age. Therefore, reduced saliency of objects in their environment may adversely affect older adults. Here, younger (6M/10F; age 22.04±3.61) and older (4M/12F; age 69.54±3.56) adults walked on four different paths, comprising straight or winding, with high or low visual contrast. We analyzed differences in lateral position (*z*_*B*_), step width (*w*), and step length (*l*). We analyzed step-to-step regulation of these within a Goal Equivalent Manifold framework. Both young and older participants exhibited substantially different stepping regulation on winding compared to straight paths. Likewise, participants in both groups exhibited more variable stepping and altered stepping regulation on low contrast compared to high contrast paths. Overall, older adults walking on winding paths took narrower steps with more variable step lengths, but exhibited similar stepping regulation. However, both stepping patterns and step-to-step regulation of older adults were significantly more disrupted by the low (compared to high) contrast paths. This was not observed in young adults. Older adults specifically prioritized step width over lateral position, likely to help maintain stability despite increased variability. Visual salience of task-relevant environmental features thus directly impacts older adults’ stepping older during walking tasks that require lateral maneuvering.

## INTRODUCTION

Falls among older (age ≥ 65 yrs) adults are extremely common (Kakara et al., 2023), contributing to $80 billion in health care costs in the US in 2020 (Haddad et al., 2024). Currently, the best predictor of a future fall is having previously fallen in the past year (Okoye et al., 2023). Visual function declines with age (Owsley, 2011; Owsley et al., 1983), and while falls are highly multi-factorial, impaired vision is an independent risk factor for falls (Black and Wood, 2005; Lord and Dayhew, 2001).

Moreover, many age-related falls occur during non-straight walking, where weight shifting is required (Robinovitch et al., 2013). This is problematic given that most real-world walking does not occur along straight lines (Glaister et al., 2007; Sannier et al., 2026), which then requires lateral stepping adjustments to navigate and avoid obstacles, etc. This in turn increases risk, as walking humans are intrinsically more unstable laterally (McAndrew et al., 2011; McAndrew Young et al., 2012; O’Connor and Kuo, 2009). Additionally, more complex real-world walking tasks also require increased visual attention (‘t Hart and Einhäuser, 2012; Matthis et al., 2018), which is likely more challenging for older adults (Berard et al., 2012; DiBianca et al., 2025; Dowiasch et al., 2015; Liu et al., 2025). Thus, gaining a better understanding of how age-related functional declines impact how people maintain balance while performing such complex real-world-like walking tasks is critical to addressing age-related balance impairments (Grosboillot et al., 2024).

From a biomechanical standpoint, we can assess how people perform such complex walking tasks by quantifying how they regulate their stepping movements at each new step (Dingwell and Cusumano, 2019; Dingwell et al., 2010). Following a Goal Equivalent Manifold (GEM) approach (Cusumano and Cesari, 2006), we can quantify how humans exploit task equifinality to correct deviations relevant to achieving specific task *goals*, while allowing more variability otherwise. For example, when humans walk on wide straight paths, they trade off correcting deviations in step widths and lateral positions relative to their path (Dingwell and Cusumano, 2019). They flexibly adapt these stepping strategies in theoretically-coherent ways when the task goals change, such as when given explicit feedback (Render et al., 2021), when walking on straight paths that slowly narrow (Kazanski et al., 2023), when they enact lateral lane shifts (Desmet et al., 2022), or when they walk on continuously winding paths (Render et al., 2025). This GEM framework thus provides an ideal framework for assessing how such stepping regulation processes change with aging (Desmet et al., 2024; Kazanski et al., 2020, 2023) and/or other locomotor impairments (Dingwell et al., 2021).

As a dynamic process, walking epitomizes “embodied” decision-making (Cisek and Green, 2024; Miller et al., 2022) as it generally requires continuous (rather than discrete) changes in direction to successfully navigate (Kessler et al., 2024). One must plan steps to negotiate obstacles and other challenges all while maintaining balance and position. To do this, humans need visual input from their external environment. Specifically, vision provides both egocentric (relation of self to objects) and allocentric (relation of objects to each other) information (Ekstrom et al., 2017; Martins et al., 2024) needed to plan stepping movements. This information, along with the desired task goal (e.g., path and/or destination, etc.), then drives motor outputs like foot placement (Domínguez-Zamora and Marigold, 2021; Drew and Marigold, 2015). Enacting these processes in real time requires people to distinguish stimuli within their environment. This is known as visual *salience*: i.e., how well a stimulus stands out from its environment (Tatler et al., 2011). Decreasing the visual salience of stepping targets degrades stepping accuracy (Domínguez-Zamora et al., 2018) and these effects are more pronounced in older adults (Black et al., 2014; Lord and Fitzpatrick, 2001).

Here, we asked young and older healthy adults to walk in four conditions, combining either straight or continuously winding paths, each presented with either high or low visual salience. We manipulated visual salience by changing the color contrast (Portengen et al., 2023) between the walking path and its background (Render et al., 2026). We hypothesized that older adults would be disproportionately challenged by continuously winding and/or low visual contrast walking paths. We tested four predictions. (1) All participants would demonstrate greater difficulty maintaining step width and/or position on winding paths than on straight paths. (2) All participants would demonstrate greater difficulty maintaining step width and/or position on low contrast paths. (3) Older adults would demonstrate greater difficulty maintaining step width and/or position than younger adults. (4) Older adults would disproportionately demonstrate greater difficulty maintaining step width and/or position on low contrast paths compared to themselves on high contrast paths, and compared to younger adults on low contrast paths.

## METHODS

This study was approved by the Institutional Review Board at the Pennsylvania State University. All participants provided written informed consent. Twenty-five young (YH; age 18-35) and thirty-one older (OH; age ≥ 65) healthy adults were screened for eligibility. Individuals were excluded if they reported taking medications, having a history of lower limb injuries or surgeries, or having musculoskeletal, cardiovascular, neurological, or visual conditions that may have compromised their gait. Individuals with blood pressure >140/90 (Frese et al., 2011), Body Mass Index (BMI) <18 or >30 kg/m^2^, or visual acuity worse than 20/40 (Hong et al., 2014) were excluded. Twenty participants in each group met all inclusion criteria and completed the study protocol. Final analyses were performed for sixteen participants in each group (Table 1) due to technical difficulties that occurred during data collection in some trials.

**Table 1.** Participant physical characteristics by age group. All values except sex are given as Mean ± Standard Deviation. P-values are from 2-sample t-tests.

| **Characteristic:** | **Young Healthy (YH):** | **Older Healthy (OH):** | **p-value:** |
|---|---|---|---|
| Sex [M/F] | 6/10 | 4/12 |  |
| Age [yrs] | 22.04 ± 3.61 | 69.54 ± 3.56 | **9.4×10<sup>-27</sup>** |
| Mass [kg] | 67.55 ± 12.80 | 70.52 ± 13.93 | 0.534 |
| Height [m] | 1.72 ± 0.07 | 1.67 ± 0.10 | 0.089 |
| BMI [kg/m<sup>2</sup>] | 22.64 ± 3.20 | 25.14 ± 3.40 | **0.040** |

Visual contrast sensitivity was assessed using the Freiburg Visual Acuity and Contrast Test (FrACT_10_, ver. 1.5; https://michaelbach.de/fract/) (Bach, 2007), which measures Weber contrast thresholds (C_Weber_) converted to logarithmic Weber contrast sensitivity scores: logCS_Weber_ = log_10_(1/C_Weber_) (Bartholomew et al., 2016; Friedel et al., 2025) (Table 2).

**Table 2.** Participant assessment scores by age group. All values are given as Mean ± Standard Deviation. P-values are from 2-sample t-tests.

| **Assessment:** | **Young Healthy (YH):** | **Older Healthy (OH):** | **p-value:** |
|---|---|---|---|
| 5mWT [m/s] | 1.322 ± 0.141 | 1.316 ± 0.139 | 0.910 |
| TUG [s] | 8.15 ± 0.87 | 9.05 ± 1.08 | **0.015** |
| FSST [s] | 7.87 ± 1.66 | 9.10 ± 1.40 | **0.030** |
| Icon-FES [/40] | 12.38 ± 1.59 | 13.38 ± 2.22 | 0.153 |
| logCS<sub>Weber</sub> [−] | 1.99 ± 0.02 | 1.85 ± 0.31 | 0.092 |

Perceived balance ability was assessed using the Iconographic Falls Efficacy Scale (Icon-FES) (Delbaere et al., 2011). Physical function was assessed from the five-meter walk test (5mWT) to estimate preferred walking speed (Ng et al., 2013), the timed-up-and-go test (TUG) (Podsiadlo and Richardson, 1991), and four-square step test (FSST) (Dite and Temple, 2002) (Table 2).

Retroreflective markers (18) were placed on each shoe at the 1^st^ and 5^th^ metatarsal heads, directly under the lateral malleolus, and on the calcaneus. Four markers were placed around the pelvis at the left and right anterior superior iliac spines (ASIS), and left and right posterior superior iliac spines (PSIS). Five markers were placed on the head: two front, two back, and one top. One final marker was placed on the right shoulder to help differentiate right and left sides of the body.

Participants walked on a 1.2 m wide treadmill (Motek Medical, Amsterdam, Netherlands; Fig. 1A) at a constant speed of 1.0 m/s while attached to an overhead safety harness. A virtual forest scene was projected onto a 240° cylindrical-shaped screen, with the speed of optic flow matched to treadmill speed.

**Figure 1.**
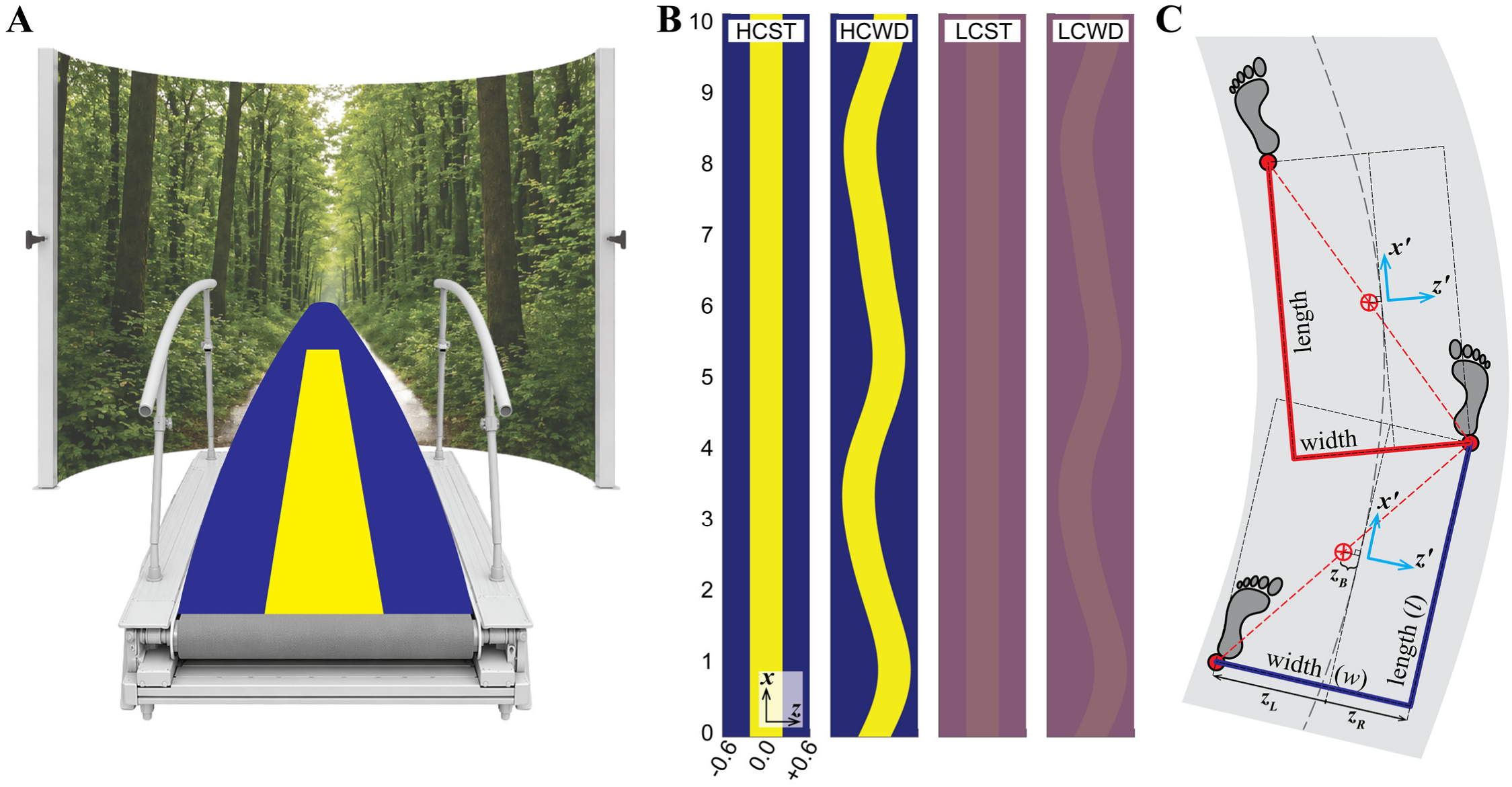
Experimental Methods and Stepping Variables. (**A**) Schematic of the Motek VR setup that participants walked in, showing approximately what they saw when walking on a high contrast straight (HCST) path. (**B**) Visual representation (plotted to scale) of a 10 m long section of each of the four paths presented: high contrast straight (HCST) and winding (HCWD) and low contrast straight (LCST) and winding (LCWD). All paths were 0.45 m wide, projected onto a 1.2 m wide treadmill belt. Winding paths (WD) were constructed using Eq. (1). High contrast (HC) conditions projected a yellow ([R,G,B] = [254,255,0]) path onto a dark blue ([R,G,B] = [0,0,129]) background. Low contrast (LC) conditions projected a slightly lighter purple ([R,G,B] = [144,102,112]) path onto a slightly darker purple ([R,G,B] = [132,86,116]) background. (**C**) Visual representation of how stepping coordinates were rotated to align with the path (Dingwell et al., 2023) for each individual step. Within these path-aligned coordinates, lateral body position was defined as *z*_*B*_ = ½(*z’*_*L*_ + *z’*_*R*_), step width was defined as *w* = (*z’*_*R*_ – *z’*_*L*_), and step length was defined as *l* = (*x’*_*Leading*_ – *x’*_*Trailing*_), depending on which foot (*L* or *R*) was leading vs. trailing for that step.

Virtual paths (Fig. 1B) were then projected onto the screen and treadmill surface (see Fig. 1A). All paths were 0.45 m wide and were either Straight (ST) or Winding (WD). Winding paths oscillated pseudo-randomly as the sum of three incommensurate sine waves (Render et al., 2024, 2025):

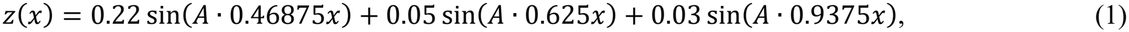

where *z*(*x*) is the lateral position of the path center, *x* is forward treadmill distance (both in meters), and *A* is a frequency scaling factor. These ST and WD paths were each projected in either High (HC) and Low (LC) visual contrast (Fig. 1B) (Render et al., 2026).

To acclimate to these tasks, each participant first completed six 90-second progressively more challenging walking trials. Participants were instructed to walk and to do their best to remain on each path for all trials. The 1^st^ acclimation trial presented a 1.2 m wide straight high contrast path. On the 2^nd^, this path was narrowed to 0.45 m wide (i.e., HCST; Fig. 1B). On the 3^rd^, this 0.45 m path was made to oscillate slowly (*A* = 1; Eq. 1). On the 4^th^, the path oscillation was increased to *A* = 3 (i.e., HCWD; Fig. 1B). On the 5^th^, the visual contrast of this path was reduced to an intermediate value between the HC and LC conditions (i.e., path [R,G,B] = [152,113,109] and background [R,G,B] = [115,66,120]). On the 6^th^ (final) acclimation trial, the winding path contrast was reduced to the full Low contrast condition (i.e., LCWD; Fig. 1B).

Following this, all participants completed three trials on each of the four test paths (Fig. 1B). They were re-instructed to do their best to remain on each path, and they again received the same auditory and visual feedback if they stepped off the path. For each path condition, participants completed one 1-minute trial, followed by two 4-minute experimental trials.

Data were recorded from the latter two trials. The order in which conditions were presented was random for each participant, but followed a fully counterbalanced Latin-square design across participants. Participants were offered rest and water as needed between trials.

Kinematic data were collected at 100 Hz using a 9-camera Vicon motion capture system (Vicon Motion Systems, Oxford, UK). Raw marker trajectories were first cleaned using Vicon Nexus software. Path position data were collected from D-Flow software (Motek Medical, Netherlands). All subsequent analyses were performed in MATLAB (MathWorks, Natick, MA). Raw marker data were filtered with a 4^th^-order Butterworth filter at a 10 Hz cutoff and then spline-interpolated to 600 Hz to more precisely identify timing of stepping events (Bohnsack-McLagan et al., 2016). Stepping events (heel strike and toe off) were identified using a marker-based algorithm (Zeni et al., 2008). For consistency, data from the last 340 steps of each trial were used to calculate all dependent measures.

When participants walked on winding paths, we re-aligned, at each step, the local coordinate system to coincide with the direction of the path at that step (Fig. 1C) (Dingwell et al., 2023; Render et al., 2025). In these local path-based coordinates, we then computed lateral position (*z*_*B*_), step width (*w*), and step length (*l*) relative to that path at that step (Fig. 1C). We then computed time series of *z*_*Bn*_, *w*_*n*_, and *l*_*n*_ for all steps *n* ∈ {1,…*N*} for the last *N* = 340 steps of each 4-minute trial. For each such time series from each trial, we then computed within-trial means, *μ*, and standard deviations, *σ*.

For each *z*_*Bn*_ and *w*_*n*_ time series, we then performed a linear error correction analysis (Dingwell and Cusumano, 2019). For each variable, *q* ∈ {*z*_*Bn*_, *w*_*n*_}, the deviation from the mean on each given step (*n*) is computed, *q*∼_n_ = *q*_n_ - *q*^*^, and then the correction made on each subsequent step, Δ*q*_(n+1)_ = *q*_(n+1)_ - *q*_n_, is plotted against *q*∼_n_ (e.g., see Figs. 3A & 5A in Results). The slope, *M*(*q*), of a linear fit to each such plot indicates the average proportion by which deviations were corrected: e.g., a slope of *M*(*q*) = −1 would indicate 100% error correction, whereas a slope of *M*(*q*) = 0 would indicate no error correction. We computed error correction slopes for each *q* ∈ {*z*_*Bn*_, *w*_*n*_} time series for every trial.

All statistical analyses were performed in Minitab (Minitab LLC, State College, PA). We treated the Straight and Winding walking paths as distinct tasks (Render et al., 2024, 2025). For each path (Straight & Winding) separately, we conducted 2-factor (Group × Contrast) mixed-effects analyses of variance (ANOVA) on each dependent measure. For standard deviation measures (σ(*z*_*Bn*_), σ(*w*_*n*_), and σ(*l*_*n*_)), we first log-transformed these data to satisfy normality and linearity assumptions of the ANOVA model. When interaction effects were found to be significant, Tukey’s pairwise post-hoc comparisons were performed to test for differences between Groups (YH vs. OH) for each path Contrast (HC vs. LC), and for differences between path Contrasts for each Group.

**Table 3.** Straight Walking Statistical Results. Differences between Groups (YH vs. OH) and path Contrasts (HC vs. LC) for the data shown in Figs. 2 & 3, including: means (*μ*) and variability (ln(*σ*)) for lateral position (*z*_*B*_), step width (*w*), and step length (*l*), and direct control slopes (*M*(*z*_*B*_) & *M*(*w*)). ANOVA results (F-statistics and p-values) are provided for main effects of Group, Contrast, and Group × Contrast interactions. Significant differences are indicated in bold.

| Data In | Variable | Group | Contrast | Group ×<br>Contrast | Tukey's<br>(Group Effects) | Tukey's<br>(Contrast Effects) |
|---|---|---|---|---|---|---|
| Fig. 2A | *μ*(*z<sub>B</sub>*) | F<sub>(1,30)</sub> = 0.20<br>p = 0.662 | F<sub>(1,94)</sub> = 2.32<br>p = 0.131 | F<sub>(1,94)</sub> = 0.11<br>p = 0.739 | N/A | N/A |
|  | *μ*(*w*) | F<sub>(1,30)</sub> = 0.27<br>p = 0.610 | F<sub>(1,94)</sub> = 1.35<br>p = 0.248 | F<sub>(1,94)</sub> = 0.08<br>p = 0.784 | N/A | N/A |
|  | *μ*(*l*) | F<sub>(1,30)</sub> = 0.15<br>p = 0.703 | F<sub>(1,94)</sub> = 1.36<br>p = 0.247 | F<sub>(1,94)</sub> = 0.65<br>p = 0.421 | N/A | N/A |
| Fig. 2B | ln(*σ*(*z<sub>B</sub>*)) | F<sub>(1,30)</sub> = 0.67<br>p = 0.418 | F<sub>(1,94)</sub> = 12.56<br>**p = 0.001** | F<sub>(1,94)</sub> = 0.14<br>p = 0.712 | N/A | N/A |
|  | ln(*σ*(*w*)) | F<sub>(1,30)</sub> = 0.03<br>p = 0.866 | F<sub>(1,94)</sub> = 3.64<br>p = 0.060 | F<sub>(1,94)</sub> = 3.24<br>p = 0.075 | N/A | N/A |
|  | ln(*σ*(*l*)) | F<sub>(1,30)</sub> = 3.89<br>p = 0.058 | F<sub>(1,94)</sub> = 1.10<br>p = 0.296 | F<sub>(1,94)</sub> = 2.42<br>p = 0.123 | N/A | N/A |
| Fig. 3B | *M*(*z<sub>B</sub>*) | F<sub>(1,30)</sub> = 5.32<br>**p = 0.028** | F<sub>(1,94)</sub> = 5.97<br>**p = 0.016** | F<sub>(1,94)</sub> = 0.31<br>p = 0.580 | N/A | N/A |
|  | *M*(*w*) | F<sub>(1,30)</sub> = 0.08<br>p = 0.780 | F<sub>(1,94)</sub> = 4.26<br>p* = 0.042 | F<sub>(1,94)</sub> = 4.83<br>**p = 0.030** | *High Contrast:*<br>YH-OH: p = 0.784<br>*Low Contrast:*<br>YH-OH: p = 0.978 | *Young:*<br>**HI-LO: p = 0.017**<br>*Older:*<br>HI-LO: p = 1.000 |

## RESULTS

### Walking on Straight Paths

Mean (*µ*) stepping parameters (Fig. 2A) were not significantly different between either Groups or Contrasts, and exhibited no significant Group × Contrast interactions (all p ≥ 0.131, Table 3).

**Figure 2.**
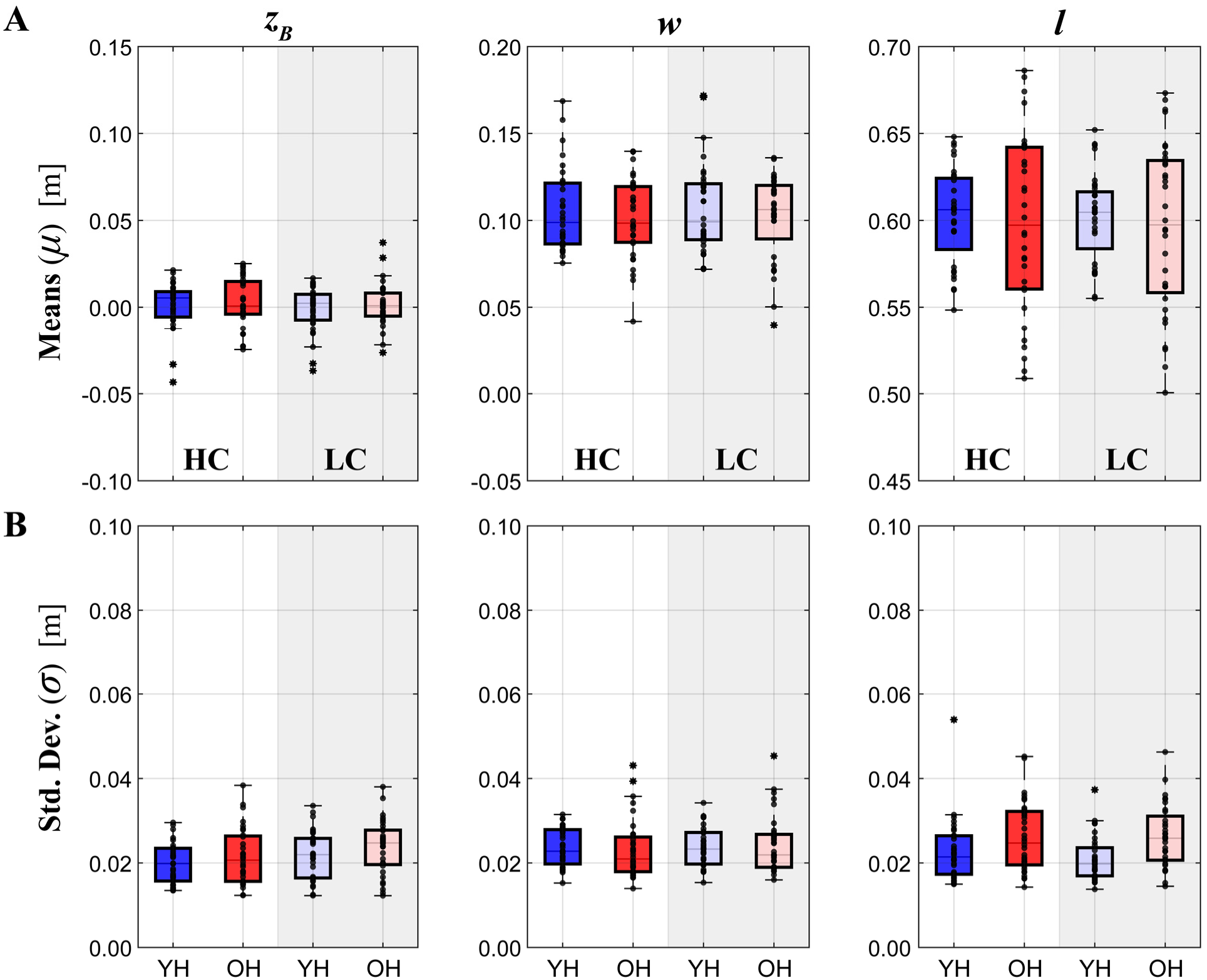
Straight Walking Stepping Parameters. (**A**) Mean values (*µ*) for lateral position (*z*_*B*_), step width (*w*), and step length, (*l*), for both Young Healthy (YH; dark and light blue) and Old Healthy (OH; dark and light red) Groups walking on both High Contrast (HC; white background) and Low Contrast (LC; gray background) paths. Plots in (**A**) have different axis limits, but are all plotted on the same scale (i.e., range). (**B**) Corresponding standard deviations (σ) for *z*_*B*_, *w*, and *l* for each Group and Contrast. Box plots display medians, 1^st^ and 3^rd^ quartiles and whiskers for 1.5 × interquartile range. Values beyond this range are indicated as asterisks. Individual trials (two trials each for each participant) are indicated by single black dots overlaid on top of the boxplots. For visual comparison, all data in both (**A**) and (**B**) are plotted on the same scales as the corresponding plots in Fig. 4. Results of all statistical comparisons are given in Table 3.

For stepping variability (Fig. 2B), both Groups exhibited significantly greater lateral position variability, ln(σ(*z*_*B*_)), on Low Contrast paths (p = 0.001, Table 3). There were no other statistically significant differences in stepping variability (all p ≥ 0.058, Table 3).

For stepping error correction (Fig. 3) on Straight paths, participants corrected step width (*w*) deviations (Fig. 3A, right) much more strongly (*M* → −1) than they corrected lateral position (*z*_*B*_) deviations (*M* → 0) (Fig. 3A, left), as expected (Dingwell and Cusumano, 2019). Both groups (YH & OH) corrected deviations in *z*_*B*_ less while walking on the Low Contrast (LC) vs. High Contrast (HC) paths (Fig. 3B, left; p = 0.016, Table 3). OH adults corrected deviations in *z*_*B*_ less compared to YH (Fig. 3B, left; p = 0.028, Table 3). YH adults corrected deviations in *w* somewhat more on the LC vs. HC paths (Fig. 3B, right; p = 0.017, Table 3). Otherwise, there were no significant differences between how YH and OH participants corrected deviations in *w*.

**Figure 3.**
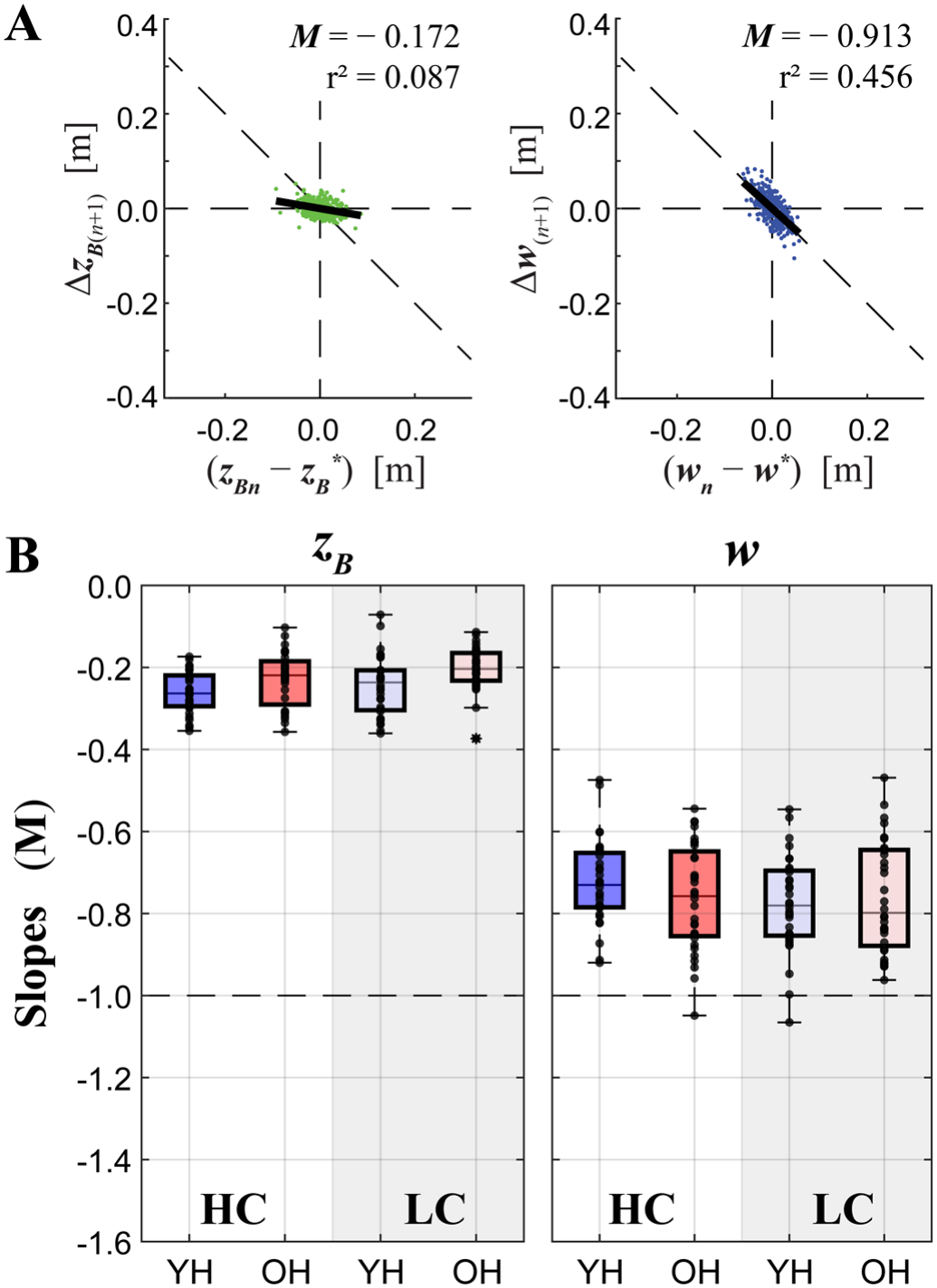
Straight Walking Linear Error Correction. (**A**) Representative linear error correction plots for *z*_*B*_ (left) and *w* (right) (Dingwell and Cusumano, 2019) from a representative trial for a representative Older (OH) participant walking on a low contrast (LC) path. At each step *n*, we plot the deviation of each variable from its target value (*z*_*B*_^*\**^ or *w*^*\**^) at that step (i.e., (*z*_*Bn*_ − *z*_*B*_ ^*\**^) or (*w*_*n*_ − *w*^*\**^)) against the corrections made on the subsequent *n*+1 step (i.e., Δ*z*_*B*(*n*+1)_ or Δ*w*_(*n*+1)_). Individual dots represent individual step pairs. Solid diagonal lines show the slopes, *M*, obtained by least-squares fit, that quantify the average degree of error corrected across all steps in that trial. A slope of *M* = −1 indicates perfect error correction, as indicated by the dashed diagonal line. (**B**) Group data for linear error correction slopes for both Young Healthy (YH) and Old Healthy (OH) Groups walking on both High Contrast (HC) and Low Contrast (LC) paths. Box plots follow the same formatting as in Fig. 2. For visual comparison, all data in both (**A**) and (**B**) are plotted on the same scales as the corresponding plots in Fig. 5. Results of all statistical comparisons are in Table 3.

### Walking on Winding Paths

On Winding paths, both groups exhibited slightly more negative (i.e., leftward) lateral positions (*μ*(*z*_*B*_)), and walked with slightly narrower mean step widths, (*μ*(*w*)), on Low (LC) vs. High (HC) contrast paths (Fig. 4A; both p < 0.0005, Table 4). Otherwise, there were no other significant main effects (all p ≥ 0.105) or interaction effects (all p ≥ 0.299) observed (Fig. 4A; Table 4).

**Table 4.** Winding Walking Statistical Results. Differences between Groups (YH vs. OH) and path Contrasts (HC vs. LC) for the data shown in Figs. 4 & 5, including: means (*μ*) and variability (ln(*σ*)) for lateral position (*z*_*B*_), step width (*w*), and step length (*l*), and direct control slopes (*M*(*z*_*B*_) & *M*(*w*)). ANOVA results (F-statistics and p-values) are provided for main effects of Group, Contrast, and Group × Contrast interactions. Where interaction effects were significant, Group and Contrast main effects should be considered unreliable. Thus, Tukey’s post-hoc comparisons are presented where relevant. Significant differences are indicated in bold.

| **Data In** | **Variable** | **Group** | **Contrast** | **Group ×<br>Contrast** | **Tukey’s<br>(Group Effects)** | **Tukey’s<br>(Contrast Effects)** |
|---|---|---|---|---|---|---|
| Fig. 4A | *μ*(*z<sub>B</sub>*) | F<sub>(1,30)</sub> = 0.48<br>p = 0.493 | F<sub>(1,94)</sub> = 23.77<br>**p < 0.0005** | F<sub>(1,94)</sub> = 1.09<br>p = 0.299 | N/A | N/A |
|  | *μ*(*w*) | F<sub>(1,30)</sub> = 1.61<br>p = 0.214 | F<sub>(1,94)</sub> = 25.92<br>**p < 0.0005** | F<sub>(1,94)</sub> = 0.28<br>p = 0.601 | N/A | N/A |
|  | *μ*(*l*) | F<sub>(1,30)</sub> = 1.92<br>p = 0.176 | F<sub>(1,94)</sub> = 2.67<br>p = 0.105 | F<sub>(1,94)</sub> = 0.05<br>p = 0.819 | N/A | N/A |
| Fig. 4B | ln(*σ*(*z<sub>B</sub>*)) | F<sub>(1,30)</sub> = 1.87<br>p = 0.181 | F<sub>(1,94)</sub> = 34.92<br>p* < 0.0005 | F<sub>(1,94)</sub> = 16.17<br>**p < 0.0005** | *High Contrast:*<br>YH-OH: p = 0.999<br>*Low Contrast:*<br>YH-OH: p = 0.070 | *Young:*<br>HI-LO: p = 0.543<br>*Older:*<br>HI-LO: **p < 0.0005** |
|  | ln(*σ*(*w*)) | F<sub>(1,30)</sub> = 1.15<br>p = 0.292 | F<sub>(1,94)</sub> = 25.27<br>p* < 0.0005 | F<sub>(1,94)</sub> = 8.63<br>**p = 0.004** | *High Contrast:*<br>YH-OH: p = 1.000<br>*Low Contrast:*<br>YH-OH: p = 0.203 | *Young:*<br>HI-LO: p = 0.455<br>*Older:*<br>HI-LO: **p < 0.0005** |
|  | ln(*σ*(*l*)) | F<sub>(1,30)</sub> = 5.76<br>**p = 0.023** | F<sub>(1,94)</sub> = 5.27<br>**p = 0.024** | F<sub>(1,94)</sub> = 0.31<br>p = 0.577 | N/A | N/A |
| Fig. 5B | *M*(*z<sub>B</sub>*) | F<sub>(1,30)</sub> = 1.56<br>p = 0.222 | F<sub>(1,94)</sub> = 9.70<br>p* = 0.002 | F<sub>(1,94)</sub> = 14.87<br>**p < 0.0005** | *High Contrast:*<br>YH-OH: p = 0.987<br>*Low Contrast:*<br>YH-OH: p = 0.161 | *Young:*<br>HI-LO: p = 0.953<br>*Older:*<br>HI-LO: **p < 0.0005** |
|  | *M*(*w*) | F<sub>(1,30)</sub> = 1.09<br>p = 0.305 | F<sub>(1,94)</sub> = 34.46<br>p* < 0.0005 | F<sub>(1,94)</sub> = 16.70<br>**p < 0.0005** | *High Contrast:*<br>YH-OH: p = 1.000<br>*Low Contrast:*<br>YH-OH: p = 0.192 | *Young:*<br>HI-LO: p = 0.590<br>*Older:*<br>HI-LO: **p < 0.0005** |

**Figure 4.**
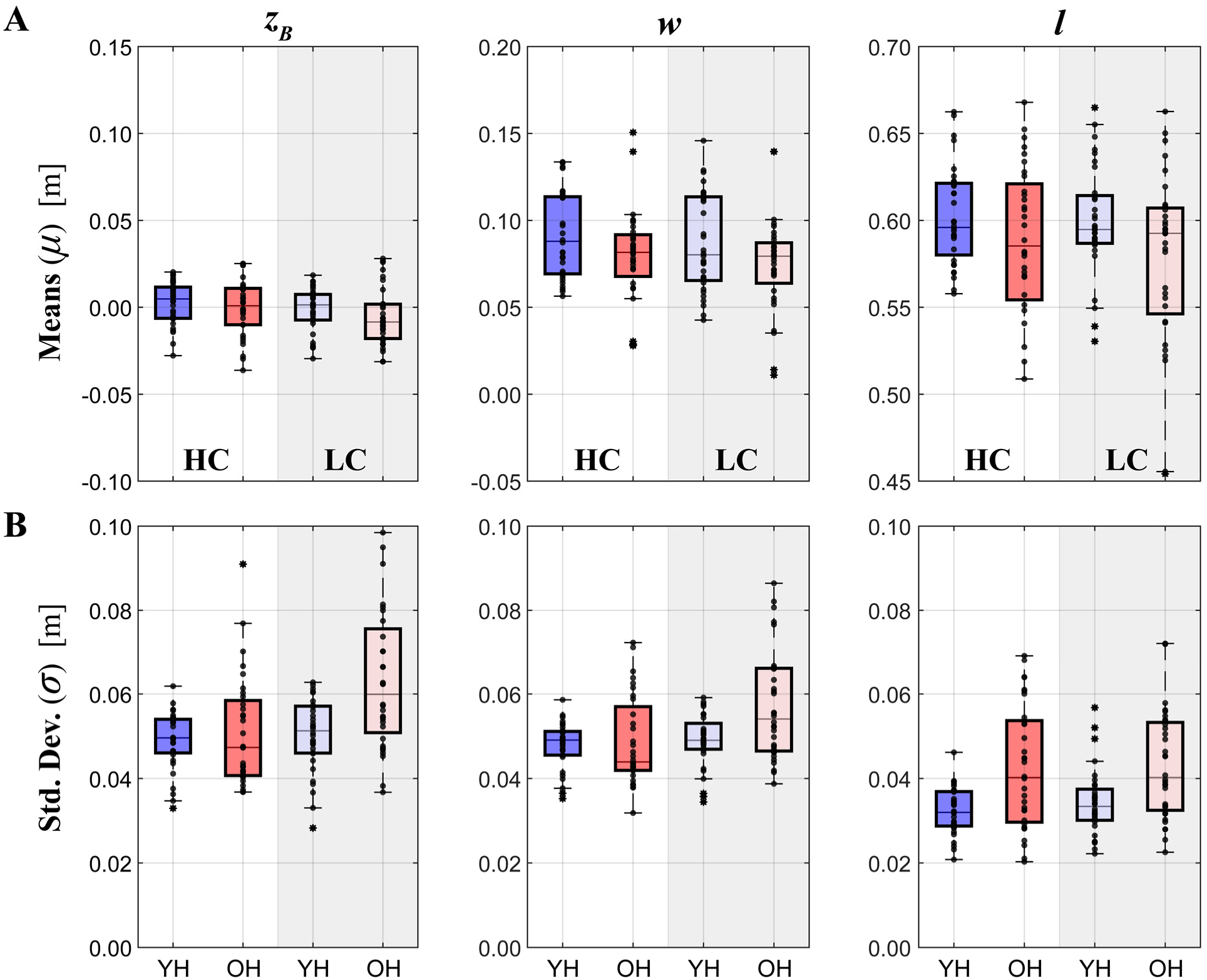
Winding Walking Stepping Parameters. **(A)** Mean values (*µ*) and **(B)** corresponding standard deviations (σ) for *z*_*B*_, *w*, and *l*, for both YH and OH Groups walking on both HC and LC paths. Box plots follow the same formatting as in Figs. 2-3. For visual comparison, all data in both (**A**) and (**B**) are plotted on the same scales as the corresponding plots in Fig. 2. Results of all statistical comparisons are given in Table 4.

For variability (Fig. 4B) of both lateral position (ln(σ(*z*_*B*_)) and step width (ln(σ(*w*)), the OH adults (but not YH adults) exhibited significantly larger increases in variability when walking on the Low Contrast (LC) compared to High Contrast (HC) paths (Fig 4B; p < 0.0005, Table 4). OH adults also exhibited significantly greater step length variability (ln(σ(*l*)) than YH adults on all paths (p = 0.023, Table 4), and both groups exhibited significantly greater σ(*l*) on LC compared to HC paths (p = 0.024, Table 4).

For step-to-step error correction (Fig. 5B), OH (but not YH) adults walking on the Low Contrast (LC) paths (only) corrected stepping errors in lateral position (*M*(*z*_*B*_)) significantly *more* and errors in step width (*M*(*w*)) significantly *less* compared to High Contrast (HC) paths (Fig 5B; both p < 0.0005, Table 4).

**Figure 5.**
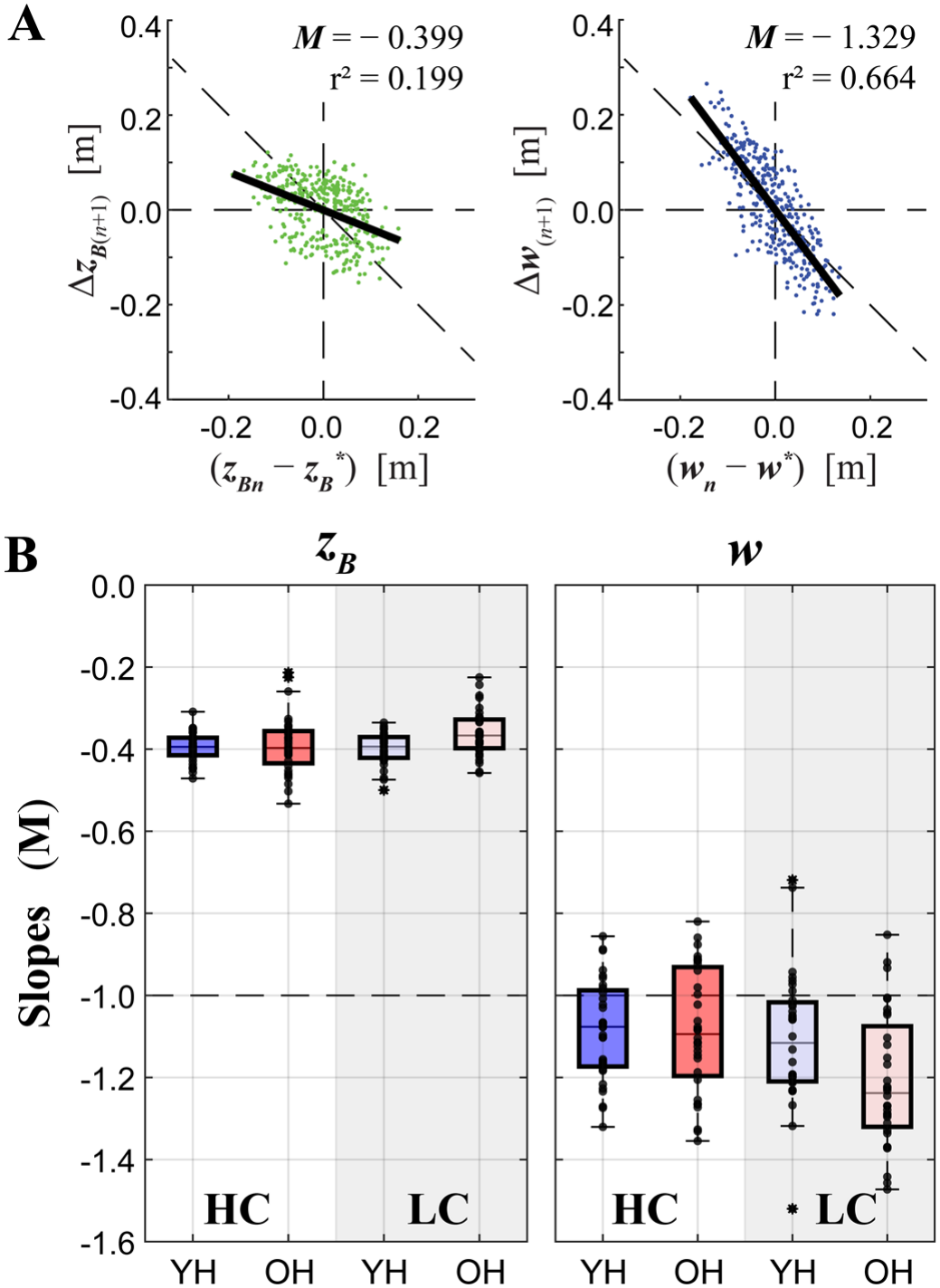
Winding Walking Linear Error Correction. **(A)** Representative linear error correction plots for *z*_*B*_ (left) and *w* (right) from the same representative OH participant as in Fig. 3A, plotted in the same manner as Fig. 3A. **(B)** Group data for linear error correction slopes for both Young Healthy (YH) and Old Healthy (OH) Groups walking on both High Contrast (HC) and Low Contrast (LC) paths. Box plots follow the same formatting as in Figs. 2-4. For visual comparison, all data in both (**A**) and (**B**) are plotted on the same scales as the corresponding plots in Fig. 3. Results of all statistical comparisons are given in Table 4.

## DISCUSSION

Falls are the leading cause of injury-related death and the rate of incidence of fatal falls is increasing (Kakara et al., 2023). Most falls occur while walking, particularly when having to make adjustments in the lateral direction (Robinovitch et al., 2013) – and everyday walking typically requires making many such adjustments (Glaister et al., 2007; Sannier et al., 2026). Thus, it is critical to consider the context of the environments in which people walk (Kessler et al., 2024; Matthis et al., 2018). Here, we quantified stepping behavior of both young and older healthy adults as they walked on straight or continuously winding (Render et al., 2025) paths. Because visual input is both critical for locomotor planning (Drew and Marigold, 2015; Martins et al., 2024), but also declines with normal aging (Dowiasch et al., 2015; Owsley, 2011), we systematically manipulated the visual contrast of the walking paths (Render et al., 2026) participants walked on.

On winding compared to straight paths, all participants exhibited substantially more variable stepping movements (Figs. 4B vs. 2B) and corrected both lateral position and step width more (Figs. 5B vs. 3B). This supports our first prediction that both young and older adults would exhibit greater difficulty maintaining step width and lateral position when walking on the winding paths. These findings were consistent with, and extend, our prior experiments using similar paths (Render et al., 2025; Render et al., 2026).

On Straight low (LC) compared to high (HC) contrast walking paths, all participants exhibited more variable lateral positions (Fig. 2B; *z*_*B*_) and corrected errors in lateral position (Fig. 3B) less. On Winding LC compared to HC walking paths, all participants also walked with narrower step widths (Fig. 4A; *w*) and more variable step lengths (Fig. 4B; σ(*l*)). These findings support our second prediction that all participants would exhibit greater difficulty navigating low contrast paths relative to otherwise-identical high contrast paths.

Compared to YH adults, OH adults walking on Straight paths corrected errors in lateral position paths less (Fig. 3B), but exhibited no differences from YH adults in variability of either lateral position or step width (Fig. 2B) or step width error correction (Fig. 3B). When walking on Winding paths, these OH adults walked with more variable step lengths (Fig. 4B), but did not exhibit significant differences from YH adults for lateral position (*z*_*B*_) or step width (*w*) variability (Fig. 4B), or for step-to-step error correction (Fig. 5B). Overall, these results do not support our third prediction that older adults would have more difficulty maintaining step width and/or position than younger adults *in general*.

However, when the OH adults walked on Low (LC) compared to High (HC) contrast paths, they exhibited significant increases in step width on Straight paths (Fig. 2B). The effects on Winding paths were more pronounced. Here, OH adults exhibited significant increases from HC to LC paths in both step width and lateral position variability (Fig. 4B; Table 4) that were not observed in YH adults. Additionally, only the OH adults corrected lateral position significantly less, and step width significantly more, on the LC compared to HC Winding paths (Fig. 5B; Table 4). YH adults did not exhibit significant HC-to-LC changes in these measures. Hence, OH adults made significantly more pronounced tradeoffs than YH adults specifically to adapt to walking on paths that were both biomechanically more challenging (winding) and simultaneously harder to see (low contrast). These findings thus strongly support both our fourth prediction and our governing hypothesis that healthy older adults would be disproportionately challenged by continuously winding and low visual contrast walking paths.

Previously, we found healthy older adults exhibited minimal changes in lateral stepping control when walking with continuous lateral perturbations (Kazanski et al., 2020), or when negotiating discrete lateral maneuvers (Desmet et al., 2024) – both in contexts where visual salience was *not* a factor. The present findings extend that work to demonstrate that, for complex walking tasks, age-related changes in visual processing (Dowiasch et al., 2015; Owsley, 2011) can negatively impact walking performance. Our work thus compliments similar findings in other walking contexts (Berard et al., 2012; DiBianca et al., 2025; Liu et al., 2025).

The OH participants we tested here were otherwise healthy adults who exhibited no declines in preferred walking speed (Table 2), a known indicator of declining physical function (Middleton et al., 2015; Studenski et al., 2011). They also expressed no increased fear of falling (Icon-FES; Table 2) (Delbaere et al., 2011). While the TUG and FSST completion times exhibited by our OH adults were significantly slower than YH (Table 2), these times remained substantially faster than cut-off values reported to indicate functional impairment for either test (Bischoff et al., 2003; de Aquino et al., 2022). Similarly, the slightly decreased visual contrast sensitivity scores of these OH adults were not significantly different from YH (Table 2) and remained well within the range reported for healthy/typical adults (Bartholomew et al., 2016; Friedel et al., 2025). The present results should be interpreted with these considerations in mind. It would be pertinent in future studies to further explore effects of visual context and biomechanical task complexity in more at-risk populations (e.g., clinical populations, more frail older adults, those with specific gait and/or visual impairments, etc.).

In conclusion, older adults regulated their stepping differently than younger adults specifically when visual contrast was low and the walking task required continuous lateral maneuvering. Specifically, they prioritized step width over lateral position, likely to help maintain stability despite being more variable. Our findings suggest these changes may be due more to degradation in perception-action coupling processes rather than the perception itself. These findings highlight the importance of considering the visual salience of the environment when investigating stepping regulation and planning in older adults during walking tasks that require continuous lateral maneuvering.

## ACKNOWLEDGEMENTS

The authors thank Dr. Anna C. Render and Ethan Friedman for their contributions and technical support throughout data collections. This work was partly supported by NIH grant 1-R01-AG049735 (to JBD).

